# A semi-quantitative analysis of lupus arthritis using contrasted high-field MRI

**DOI:** 10.1101/218503

**Authors:** Eric S Zollars, Russell Chapin

**Author notes:** Corresponding author: Eric Zollars, Fax: Address: 96 Jonathan Lucas ST, MSC 632, MUSC, Charleston, SC 29466. No commercial support or conflicts of interest.

## Abstract

**Objective:** Arthritis in systemic lupus erythematosus is poorly described and there is no objective measure for quantification of the arthritis. We aim to develop MRI as a research tool for the quantification of lupus arthritis.

**Methods:** Patients were eligible for entry into the study if they were evaluated at the MUSC Lupus Center and determined by their treating physician to have active hand arthritis due to SLE. Standard of care lupus activity measures were collected along with a detailed physical exam. MRI images were obtained using standard musculoskeletal sequences with gadolinium contrast. Semi-quantitative scoring of the images used the OMERACT RAMRIS system.

**Results:** RAMRIS demonstrates large amounts of synovitis, tenosynovitis, bone marrow edema and erosive disease in only a minority of patients. Some patients were not scored as having any synovitis or tenosynovits. We describe potential features of lupus arthritis that are not captured in the RAMRIS scores and may be contributing to symptoms.

**Conclusion:** Lupus arthritis is an entity separate from rheumatoid arthritis and requires the development of new quantitative methods to describe it. MRI findings suggest the inadequacy of a typical lupus musculoskeletal measures and even swollen/tender joint counts to assess the level of disease activity.

## Introduction

Systemic lupus erythematosus (lupus) is an autoimmune disease that can involve any organ system. While kidney and nervous system involvement are the more severe manifestations, the involvement of the skin and the musculoskeletal systems are the most common manifestations. The myriad rashes of lupus are well-described with the recent SLICC criteria describing 12 separate sub-types^1^. However, the musculoskeletal manifestations of lupus described by this criteria include only synovitis of at least two joints and tenderness of two joints with 30 minutes of morning stiffness. Clinical experience suggests that this an inadequate representation of the myriad symptoms experienced by lupus patients with true inflammatory arthritis.

Improved description and quantification of lupus arthritis is necessary to move lupus treatment into an era of precision medicine. First and foremost, the symptoms of lupus arthritis are experienced by 90% of patients and most patients consider it one the most disabling features of the disease^2^. Second, most lupus patients entered into interventional clinical trials are required to have moderate disease activity, frequently one component is arthritis. Unfortunately, the quantitative measures we currently have to describe lupus arthritis such as SLEDAI-2K and BILAG are rough measures and not particularly responsive to change. This has led to modifications such as the SLEDAI-2000 Responder Index 50 (SRI-50) and the BILAG-Based Composite Lupus Assessment (BICLA) to detect partial responses. A comparison of these measures in clinical practice is discussed in Thanou et al.^3^. The complexity of SLE necessitates that these indices attempt to account for all the possible changes in disease activity in multiple organ systems. For the most part, they do so and correspond to physicians’ global assessment of disease activity and intention to treat. Nevertheless, within the musculoskeletal system, most measures continue to use mostly subjective assessments of disease activity and improvement. A third reason why improved quantification is necessary is the general difficulty of assessing lupus arthritis on physical exam in the clinical setting. In the vast majority of cases lupus arthritis patients do not have the easily palpable synovial hypertrophy of rheumatoid arthritis. The obesity epidemic has not spared the lupus population and significant adipose tissue can mask subtle swelling. Further, as fibromyalgia coexists with SLE in approximately 20% of patients^4^ the subjective measures of tenderness and stiffness become even less reliable.

The utility of advanced imaging is just beginning to be applied to lupus arthritis. In 2003, early investigations with MRI noted the different features of SLE arthritis compared to RA^5^ particularly the presence of edematous tenosynovitis and capsular swelling. Since that time, a few new evaluations of lupus arthritis using MRI were reported^6–8^ with most of the focus on developing advanced imaging for RA. Recently, however, Mosca and colleagues evaluated a series of lupus patients with both ultrasound and MRI^9^. They found that approximately half of patients judged to be clinically quiescent on clinical exam had inflammation when evaluated with one of these imaging techniques. Additionally, they were able to show a significant amount of erosive damage that was not detected by standard diagnostic radiology. They hypothesize that these ‘sub-clinical’ findings are the causes of subsequent damage and warrant therapeutic intervention. Ball and colleagues investigated active lupus arthralgia with contrasted high-field MRI and found a large majority of patients with at least low-grade synovitis and erosive disease.

In rheumatoid arthritis, there are validated measures of arthritis using advanced imaging (MRI and ultrasound) in this disease. The Outcome Measures in Rheumatology (OMERACT) Rheumatoid Arthritis MRI Scoring System (RAMRIS) is a validated measure of the inflammatory features of RA seen on MRI^10^. In multiple clinical trials^11,12^ MRI is a reliable, responsive and early indicator of outcome. The increased objectivity and quantitation of the measure can allow for quite small trials that have power to show response. This was in as few as 27 patients in Ostergaard et al.^13^ The RAMRIS is composed of scores for erosions, bone marrow edema and synovitis. It has also been supplemented with a score for tenosynovitis^14^. The combination of synovitis, tenosynovitis and bone marrow edema is used to indicate a total MRI inflammatory score and is lower in patients in clinical remission^15^.

This study assessed the utility of RAMRIS in scoring lupus arthritis as well as identify features of lupus arthritis that are incompletely captured by the RAMRIS.

## Patients and Methods

Twenty lupus patients were recruited to this study from within the MUSC rheumatology practice. Approximately 1200 SLE patients are seen annually at MUSC about 2/3 are self-identified as African American. The entry criteria for this study was simply that the patient have SLE by ACR criteria and that the treating rheumatologist believed that the patient had significant inflammatory musculoskeletal lupus activity (defined loosely as lupus arthritis). The BILAG ‘intention to treat or increase treatment’ was used as a guide, i.e. the current lupus manifestations are enough to justify initiating or increasing therapy for lupus activity (whether or not that was done). Notably, the patients were not entered into the study to evaluate IF they had presence of inflammatory activity—the confidence of the treating rheumatologist was high that the patient did have arthritis.

### Clinical scoring of arthritis

There are a variety of methods to score arthritis in lupus. While SLEDAI is collected as standard-of-care we focused on a swollen and tender joint count. A single evaluator (EZ) scored each of the MCP and PIP joints and the wrist as swollen, tender or both (maximum score of 9 for each). Further record was made of tenderness in the forearm, dorsal hand, palm tenderness proximal to MCPs and pain with flexion/extension of the wrist and fingers. The rest of the routine clinical data was collected during the visit (presence of activity in other organ systems). A standard continuous measure of disease activity on a scale of 0-3 scored on a visual analogue scale^16^ was collected as well. Standard clinical practice involves assessing a patient-reported pain scale from 0-10 on a visual analogue scale.

### MRI scoring of arthritis

Patients who consented to the study had an MRI scan of the wrists and hands performed on the same day. The MRI machine was a whole-body Siemens 3T machine using a medium 4 channel flex coil. Standard sequences including T1 (axial, coronal), T2, Proton Density Fat Saturated, STIR (short-tau inversion recovery) and four post-contrast were used. The images were scored by Dr. Russell Chapin (MSK radiologist) and Eric Zollars (rheumatologist). The scoring method was the (Outcome Measures in Rheumatology Clinical Trials (OMERACT)) for assessing activity in rheumatoid arthritis, RAMRIS^10^. It includes scores for synovitis, bone edema and erosions. Subsequent work led to the development of a score for tenosynovitis^14^. The RAMRIS scores erosions (23 anatomic areas, range 0-10), synovitis (7 anatomic areas, range 0-3), and bone marrow edema (23 anatomic areas, range 0-3). The tenosynovitis score is scored in 10 areas with a score of 0-3. Thus, there is a maximum score of 230 for erosions, 21 for synovitis, 69 for bone marrow edema and 30 for tenosynovitis. Tenosynovitis is scored only in the wrist while the other scores are for the MCPs as well as the wrist. The images were scored independently by the two evaluators without access to any patient information. After 10 patients were scored we reviewed the images together to arrive at consensus scores. This was done twice.

### Statistics

Descriptive statistics are presented as mean ± standard deviation and N (%) for continuous and categorical variables, respectively. To assess the relationships between self-reported pain with RAMRIS component scores, and between swelling/tenderness scores with RAMRIS scores, Spearman’s rank correlation was used. To assess the consistency of scoring between raters for each of the RAMRIS scores, an intra-class correlation coefficient was calculated. All analyses were performed using SAS v9.4. Statistical significance was assessed at α = 0.05. Given the exploratory nature of the study, no correction for multiple comparisons was made.

## Results

A total of twenty patients were enrolled (Table 1), of whom 75% (n=15) were African Americans. The average age was 41 ± 16.3 years. Nearly all patients were on immunosuppression in addition to hydroxychloroquine. Treatments for the patients are in Table S1. The exceptions to immunosuppression were patients 4 and 20 who were new diagnoses and patient 18 who was in between stopping one agent and starting another. The majority of patients did not have serologic markers of lupus disease activity. Ant-dsDNA was measured using ELISA. If a positive is considered above the manufacturer’s range then 8 of 20 had a positive value. If the SLICC criteria of two times upper limit of normal is used than only 4 of 20 had a positive anti-dsDNA. Six of 19 (32%) had hypocomplementemia, with only one patient having an isolated low C4. Jaccoud’s was clinically apparent in three patients all of whom were in their 30s. Concomitant fibromyalgia was present in four of 20 patients. Other than the serologic markers of disease activity, seven patients had lupus manifestations that would have been counted by SLEDAI. These were leukopenia in three (attributed to lupus rather than immunosuppression), five with rash or mucocutaneous and only one patient with concomitant nephritis. No neurologic manifestations were observed as active. The patients’ historical disease associations are shown in Table S1 as well. No patient had prior severe manifestations of SLE (other than the one patient with on-going nephritis). Four patients had previous episodes of serositis. Three patients had previously had episodes of thrombocytopenia.

Table 1

Table 1
|  |  |
| --- | --- |
| **Age** | 41±16 |
| **BMI** | 31±8 |
| **Ethnicity** | 75% |
| African American | 10% |
| Hispanic | 5% |
| White | 5% |
| Asian |  |
| **Jaccoud's** | 15% |
| **Lupus activity markers** |  |
| Elevated anti-dsDNA | 40% |
| 2x ULN elevated anti-dsDNA | 20% |
| Hypocomplementemia | 32% |

We compared the MRI scores of the radiologist (Chapin) and the rheumatologist (Zollars) prior to arriving at the consensus scores. There were very few areas of significant disagreement when looking at every joint. Differences tended to be by small amounts (i.e. scoring an erosion as 2 out of 10 or 3 out of 10). Higher variability was seen in scoring bone marrow edema due to subtle variations among low values. An intra-class coefficient was calculated^17^ and was 0.956 for the erosion score, 0.907 for BME, 0.969 for synovitis and 0.936 for tenosynovitis. These scores are quite high and show both the ease of application of the atlas-based RAMRIS scores as well as our internal consistency.

The MRI scores are shown in Table 2. The large majority (90%) had some detectable erosions. One patient with long-standing disease and a positive rheumatoid factor had significant erosive disease (erosion score 38). A majority (55%) of patients had some detectable bone marrow edema, however only two patients had high signal. As was expected, 60% of patients had some RAMRIS synovitis, though most was low grade. Tenosynovitis at the wrist was also seen in a majority of patients (85%). A correlation of 0.6 was seen between synovitis and tenosynovitis scores, however there were patients with a larger component of tenosynovitis compared to synovitis (right panel in Figure 1).

Table 2

Table 2
| Patient | Erosion | BME | Tenosynovitis | Synovitis | Pain | RF | Swollen | Tender | FM |
| --- | --- | --- | --- | --- | --- | --- | --- | --- | --- |
| 1 | 0 | 0 | 5 | 1 | 2 | neg | 3 | 2 | N |
| 2 | 38 | 14 | 3 | 1 | 6 | pos | 4 | 5 | N |
| 3 | 8 | 2 | 0 | 2 | 8 | ND | 1 | 7 | Y |
| 4 | 4 | 5 | 7 | 11 | 3 | ND | 5 | 0 | N |
| 5 | 4 | 1 | 0 | 0 | 9 | neg | 6 | 2 | N |
| 6 | 2 | 0 | 0 | 1 | 6 | neg | 2 | 4 | Y |
| 7 | 1 | 1 | 0 | 1 | 0 | ND | 3 | 5 | N |
| 8 | 2 | 6 | 9 | 1 | 0 | neg | 7 | 8 | N |
| 9 | 9 | 1 | 10 | 15 | 5 | pos | 6 | 3 | N |
| 10 | 2 | 36 | 6 | 3 | 3 | ND | 3 | 3 | N |
| 11 | 4 | 4 | 0 | 4 | 7 | ND | 4 | 9 | N |
| 12 | 2 | 0 | 1 | 2 | 5 | pos | 5 | 5 | N |
| 13 | 0 | 0 | 0 | 0 | 10 | pos | 0 | 1 | Y |
| 14 | 18 | 0 | 17 | 17 | 9 | neg | 6 | 7 | N |
| 15 | 0 | 0 | 0 | 0 | 3.5 | neg | 4 | 8 | N |
| 16 | 1 | 0 | 0 | 7 | 10 | ND | 2 | 6 | N |
| 17 | 1 | 0 | 7 | 10 | 7 | neg | 7 | 7 | N |
| 18 | 9 | 0 | 1 | 1 | 7.5 | neg | 2 | 8 | Y |
| 19 | 10 | 6 | 8 | 1 | 1 | neg | 0 | 1 | N |
| 20 | 12 | 36 | 15 | 3 | 1 | pos | 8 | 4 | N |

RF: rheumatoid factor, ND: not done. FM: presence of fibromyalgia.

RF: rheumatoid factor, ND: not done. FM: presence of fibromyalgia.

**Figure 1.**
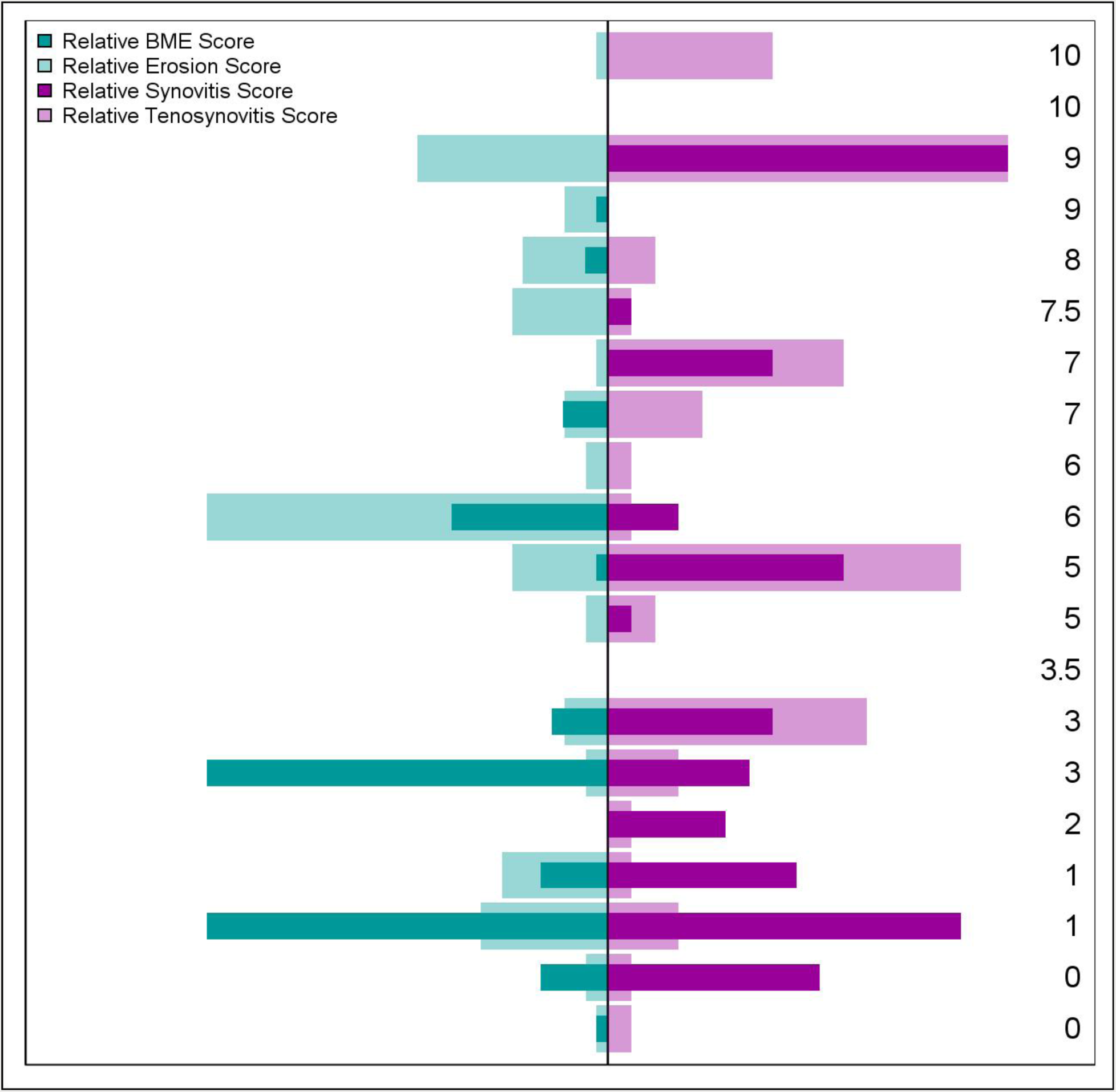
This shows the distribution of the RAMRIS component scores. The patient with the highest pain score is at the top and the lowest pain score at the bottom. The scores are normalized. Erosion and bone marrow edema are in the left panel. Synovitis and tenosynovitis are in the right panel.

An impetus of the study is develop an objective, quantitative measure of lupus arthritis. This includes attempts to uncover manifestations of joint involvement by MRI that are below the sensitivity of the physical exam. Figure 1A shows the low association between swollen joints detected by exam and those reported as tender by the patient. There is a poor association between swelling over the joints and patient reported tenderness. There are patients with significant swelling/minimal tenderness as well as those with minimal swelling/significant tenderness. Of note, the four fibromyalgia patients have the fewest number of swollen joints (0-2) with higher pain scores (6-10) but have varied tenderness scores (1-8). Comparing the objective measure of joint swelling with the various measures of inflammation on MRI shows the best correlation with synovitis (R=0.57) (Figure 1c), moderate correlation with tenosynovitis (R=0.42) (Figure 1d) and low correlation with bone marrow edema (R=0.24) (Figure 1b). The total inflammatory score (the sum of synovitis, tenosynovitis and bone marrow edema) has an intermediate correlation of 0.50 (not shown) with swelling. The stronger correlation between swelling and synovitis is not surprising as the swollen/tender joint count is based on rheumatoid arthritis, a disease with predominant synovitis. Adding just the synovitis and tenosynovitis scores together did not increase the correlation with the swelling count.

Associations with patient’s pain scores and the various features of the MRI score are shown in Figures 1E-1H). We had hypothesized that the amount of damage assessed by MRI would associate more strongly with a patient’s pain score. The association was weak (R=0.1) (Figure 1E). The strongest association with pain was the tenosynovitis score, but this too was weak (R=0.22). Synovitis and bone marrow edema did not have significant associations with patient pain scores. Notably, the two patients with severe bone marrow edema had minimal pain scores of three and one.

We conducted an extensive evaluation of the images to uncover other features of inflammatory arthritis not scored by RAMRIS (Figure 3). Every series was annotated with potentially inflammatory features. While multiple findings were identified in individual patients there were some common features seen in multiple patients. Focusing on those patients who has synovitis scores of zero but the presence of swollen joints on exam revealed joint effusions that neither enhanced with gadolinium nor had thickened, proliferative synovium. These are appropriately scored as zero by the RAMRIS criteria but are obviously abnormal on T2 MRI signals. The patients hand was kept in a neutral position during the scan so often the effusion was seen more in the volar recess. Another group of abnormalities seen on MRI was increased fluid signal around the flexor tendons between the MCPs and PIPs, likely representing a tenosynovitis. In the three examples shown in Figure 3B they all had notable signal in the fingers with minimal tenosynovitis scores in the wrist. Finally, Figure 3C shows patients with tendon effusions proximal to MCPs. This was seen in multiple patients, some of whom also had significant tenosynovitis in the wrist (unlike the above MCP-PIP tendon effusions). When we started noticing these findings we began assessing tenderness in the palm proximal to the MCPs and were able to note some association.

**Figure 2.**
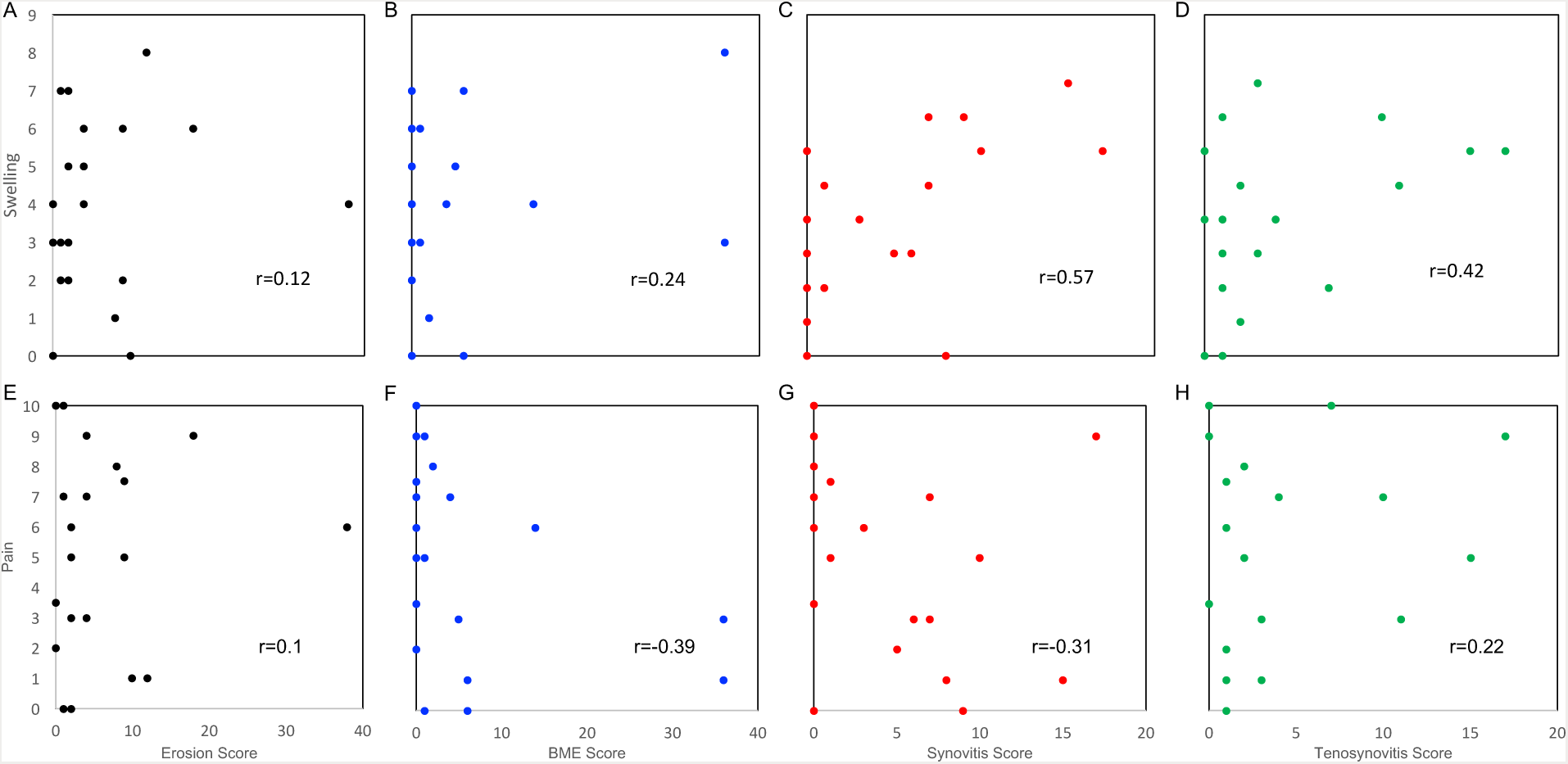
The first row shows the association between evaluator assessed swelling and the RAMRIS component scores. A: erosion, B: bone marrow edema, C: synovitis, D: tenosynovitis. The second row shows the association with patient reported pain and the RAMRIS component scores. E: erosion, F bone marrow edema, G: synovitis and H: tenosynovitis. Pearson’s correlation, r, is reported in the inset.

**Figure 3.**
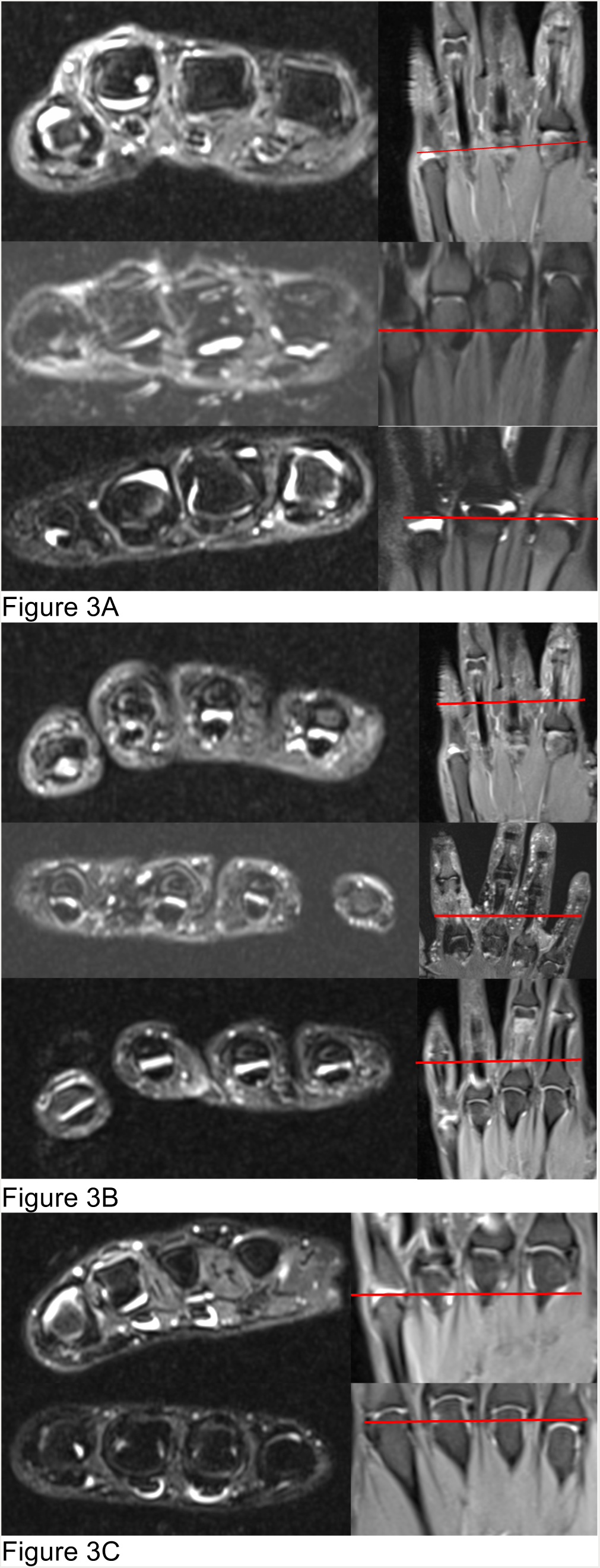
The left side of each image is a T2 axial cut on the left showing the relevant finding and the image on the right is a T2 image with a red bar to show where the cut was taken. (A). These images show non-enhancing joint effusions on three patients (5,15,16) who all have synovitis scores of 0 but swollen joints on physical exam. (B) Show flexor tendon effusions between the MCP and PIP on three patients (5, 7, 8) who have tenosynovitis scores of 0, 1 and 1 at the wrist. (C) These images show flexor tendon effusions in the palm proximal to the MCPs in two patients (8, 11) who have tenosynovitis scores of 1 and 4 at the wrist.

## Discussion

The goal of this study was to begin developing a method to quantitate active lupus arthritis. We have started with the RAMRIS semi-quantitative method that has years of development and validation within rheumatoid arthritis. The long-term goal is to develop a quantitative index that is responsive to change to allow small, focused interventional trials in lupus arthritis.

All patients in this study were thought by the treating rheumatologist to have active disease. These would be similar to those patients considered for a clinical trial. Though notably, with current typical criteria in SLE clinical trials (typically a SLEDAI of 6, 4 points that must come from non-laboratory criteria) only 40% would have met criteria for inclusion. Two of three of the patients with the most active arthritis by RAMRIS would not meet criteria due to receiving only the four points for SLEDAI arthritis. Two of the four patients with the most active tenosynovitis by RAMRIS would not meet criteria for entry. Of the two patients with profound bone marrow edema, a process in RA that is thought to indicate progression into destructive arthritis, only one would have met entry criteria.

In a large active cohort of lupus patients, those patients referred for the study had relatively mild disease outside of the musculoskeletal domain. Granted, significant nephritis or CNS disease was an exclusion but this is common in SLE trial design. This has indications for trial design in lupus arthritis. An interventional trial targeted to lupus arthritis should be able to recruit significant numbers of patients who do not have significant extra-articular disease. Then, methods to reduce heterogeneity such as removing background medications^18^ can be performed safely. Further, with an objective, quantitative marker we should have more sensitivity to detect response.

Heterogeneity was seen in the MRI measures of lupus arthritis and neither patient reported pain/ tenderness nor lupus activity serologies were associated with these measures. The patients with the highest synovitis scores (patients 14, 9), highest tenosynovitis scores (patients 14, 20) and highest bone marrow edema scores (patients 10, 20) all had normal lupus activity serologies. As shown in Figure 2 individual patients could have different quantities of erosions, synovitis, tenosynovitis and bone marrow edema.

Mild erosive damage was seen in many patients and some had severe RA-like damage. The very low scores are of unclear clinical significance. Low-grade erosions were reported in 2.2% of MCP joints and 1.7% of wrist joint bones^19^ in a study applying RAMRIS to healthy controls. More recently, Tani and colleagues applied RAMRIS to RA, SLE arthritis and healthy controls using non-contrasted low-field extremity dedicated MRI^20^. They were focused on damage and found low RAMRIS scores in many wrists of healthy controls. They found average scores of 5±2.9, 9.1±8.6, 13.4±9.4 in healthy controls, lupus arthritis and rheumatoid arthritis. By a similar metric our patients have a mean erosion score of 6.3±8.9, showing a slightly lower burden of erosive disease but similar variability. They proposed a ‘normality’ threshold above which only 2.5% of healthy controls would be expected to score. With this metric 63% of their RA patients and 24% of their SLE patients surpassed the normality threshold. Three of our 20 (15%) patients meet this criteria. The variability even among these three patients is worth commenting upon. Patient 2 (erosion score 38) has long-standing disease, elevated bone marrow edema score and a positive rheumatoid factor. She had minimal synovitis and tenosynovitis. Patient 14 (erosion score 18) has no bone marrow edema and a negative rheumatoid factor. She had the highest synovitis and tenosynovitis scores in this cohort. Patient 20 (erosion score 12) was a new diagnosis, very high bone marrow edema and a positive rheumatoid factor. She had a very high synovitis score as well. It is unclear based on the MRI findings from the patients in this cohort who would fit the definition of ‘rhupus’, a loose term indicating an overlap of rheumatoid arthritis and lupus.

There is no ‘gold standard’ for the quantification of lupus arthritis thus we have nothing to compare with the performance of RAMRIS other than clinical exam and typical metrics in interventional clinical trials. To this end we compared patients who had objective findings of swelling on exam and low RAMRIS component scores. We found joint effusions that would not be scored as there was no enhancement with gadolinium and no synovial proliferation. We note also the tenosynovitis throughout the hand and fingers seen in these SLE patients even when tenosynovitis was minimal in the wrist. It is worth recalling that RAMRIS is the *Rheumatoid Arthritis* MRI Scoring system and perhaps we will need an SLE MRI Scoring system (SLEMRIS).

At this time we cannot comment on response to change as we have not reported any longitudinal follow up. Thus, we do not know the features that are most likely to have response to typical therapies. It will be interesting to see if RA-like synovitis is responsive to typical RA treatments while perhaps more SLE-like findings do not. Nearly all of these patients were on immunosuppression in addition to hydroxychloroquine *with* the persistence of findings described here. This finding alone suggests the need for more specific therapies for lupus arthritis. MRI, though expensive, if shown to be responsive to clinical change may have utility in a research setting for the assessment and monitoring of lupus arthritis.

## Supplemental

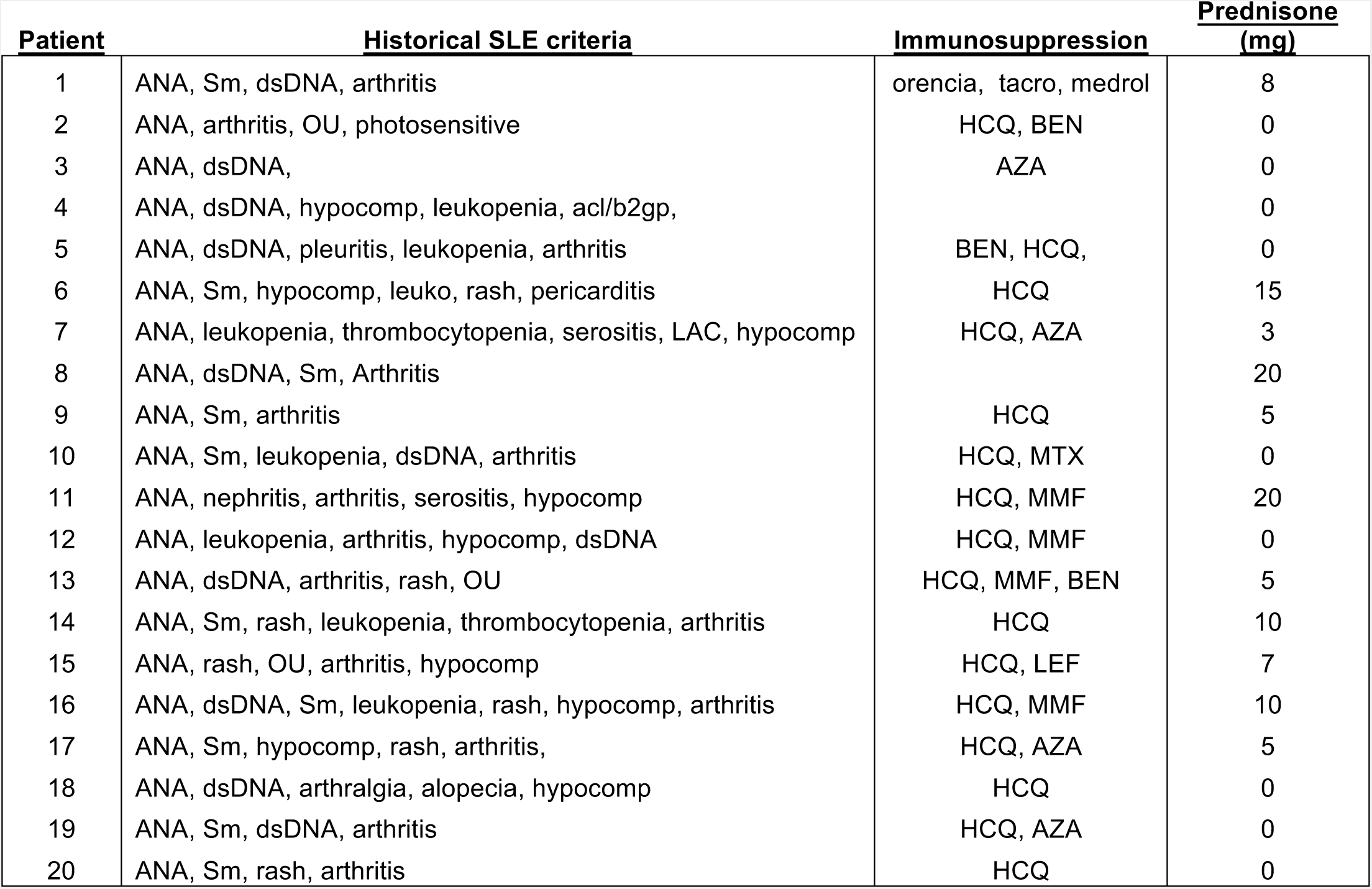

OU: oral ulcers; hypocomp: hypocomplementemia; acl/b2gp: antiphospholipid antibodies; leuko: leukopenia (or lymphopenia); LAC: lupus anticoagulant; Sm: anti-smith; tacro: tacrolimus; HCQ: hydroxychloroquine; BEN: benlysta; AZA: azathioprine; MTX: methotrexate; MMF: mycophenolate; LEF: leflunomide.

## References

1. Petri, M. et al. Derivation and validation of the Systemic Lupus International Collaborating Clinics classification criteria for systemic lupus erythematosus. Arthritis Rheum. 64, 2677–2686 (2012).

2. Waldheim, E. et al. Extent and characteristics of self-reported pain in patients with systemic lupus erythematosus. Lupus 22, 136–143 (2013).

3. Thanou, A., Chakravarty, E., James, J. A. & Merrill, J. T. Which outcome measures in SLE clinical trials best reflect medical judgment? Lupus Sci. Med. 1, e000005 (2014).

4. Middleton, G. D., McFarlin, J. E. & Lipsky, P. E. The prevalence and clinical impact of fibromyalgia in systemic lupus erythematosus. Arthritis Rheum. 37, 1181–1188 (1994).

5. Ostendorf, B., Scherer, A., Specker, C., Mödder, U. & Schneider, M. Jaccoud’s arthropathy in systemic lupus erythematosus: Differentiation of deforming and erosive patterns by magnetic resonance imaging. Arthritis Rheum. 48, 157–165 (2003).

6. Sá Ribeiro, D. et al. Magnetic resonance imaging of Jaccoud’s arthropathy in systemic lupus erythematosus. Joint Bone Spine 77, 241–245 (2010).

7. Boutry, N., Hachulla, É., Flipo, R.-M., Cortet, B. & Cotten, A. MR Imaging Findings in Hands in Early Rheumatoid Arthritis: Comparison with Those in Systemic Lupus Erythematosus and Primary Sjögren Syndrome. Radiology 236, 593–600 (2005).

8. Ball, E. M. A. et al. A study of erosive phenotypes in lupus arthritis using magnetic resonance imaging and anti-citrullinated protein antibody, anti-RA33 and RF autoantibody status. Rheumatol. Oxf. Engl. 53, 1835–1843 (2014).

9. Mosca, M. et al. The role of imaging in the evaluation of joint involvement in 102 consecutive patients with systemic lupus erythematosus. Autoimmun. Rev. doi:10.1016/j.autrev.2014.08.007

10. Østergaard, M. et al. An introduction to the EULAR–OMERACT rheumatoid arthritis MRI reference image atlas. Ann. Rheum. Dis. 64, i3–i7 (2005).

11. Conaghan, P. G. et al. Assessment by MRI of inflammation and damage in rheumatoid arthritis patients with methotrexate inadequate response receiving golimumab: results of the GO-FORWARD trial. Ann. Rheum. Dis. 70, 1968–1974 (2011).

12. Peterfy, C. et al. MRI assessment of suppression of structural damage in patients with rheumatoid arthritis receiving rituximab: results from the randomised, placebo-controlled, double-blind RA-SCORE study. Ann. Rheum. Dis. annrheumdis-2014-206015 (2014). doi:10.1136/annrheumdis-2014-206015

13. Østergaard, M. et al. MRI assessment of early response to certolizumab pegol in rheumatoid arthritis: a randomised, double-blind, placebo-controlled phase IIIb study applying MRI at weeks 0, 1, 2, 4, 8 and 16. Ann. Rheum. Dis. annrheumdis-2014-206359 (2014). doi:f10.1136/annrheumdis-2014-206359

14. Haavardsholm, E. A., Østergaard, M., Ejbjerg, B. J., Kvan, N. P. & Kvien, T. K. Introduction of a novel magnetic resonance imaging tenosynovitis score for rheumatoid arthritis: reliability in a multireader longitudinal study. Ann. Rheum. Dis. 66, 1216–1220 (2007).

15. Ranganath, V. K. et al. Comprehensive appraisal of magnetic resonance imaging findings in sustained rheumatoid arthritis remission: a substudy. Arthritis Care Res. 67, 929–939 (2015).

16. Petri, M., Hellmann, D. & Hochberg, M. Validity and reliability of lupus activity measures in the routine clinic setting. J. Rheumatol. 19, 53–59 (1992).

17. Rosner, B. Fundamentals of Biostatistics. (Cengage Learning, 2010).

18. Merrill, J. T. et al. The Biomarkers of Lupus Disease Study: A Bold Approach May Mitigate Interference of Background Immunosuppressants in Clinical Trials. Arthritis Rheumatol. Hoboken NJ 69, 1257–1266 (2017).

19. Ejbjerg, B. et al. Magnetic resonance imaging of wrist and finger joints in healthy subjects occasionally shows changes resembling erosions and synovitis as seen in rheumatoid arthritis. Arthritis Rheum. 50, 1097–1106 (2004).

20. Tani, C. et al. MRI pattern of arthritis in systemic lupus erythematosus: a comparative study with rheumatoid arthritis and healthy subjects. Skeletal Radiol. 44, 261–266 (2015).

